## Supplementary figures and images for "MEANtools: multi-omics integration towards metabolite anticipation and biosynthetic pathway prediction"

### Supplemental Figure 1

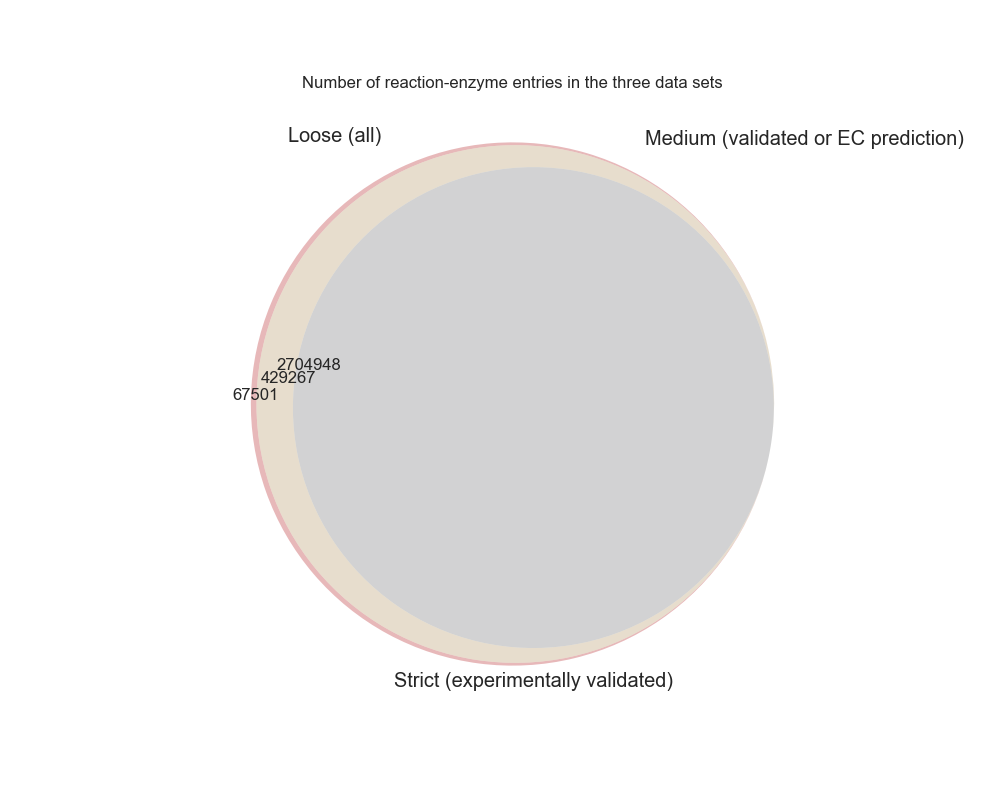

### Supplemental Figure 2

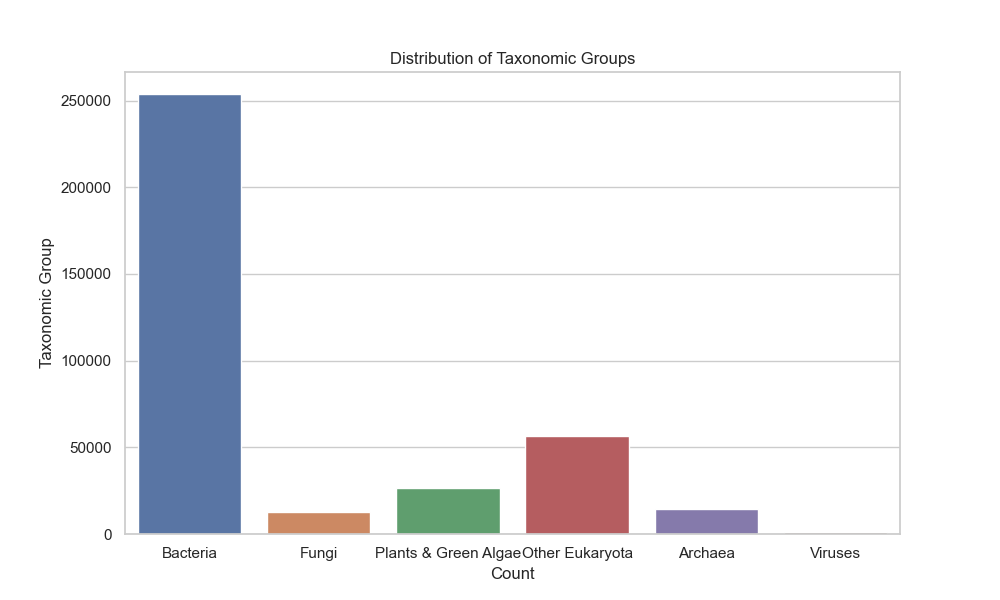

### Supplemental Figure 3

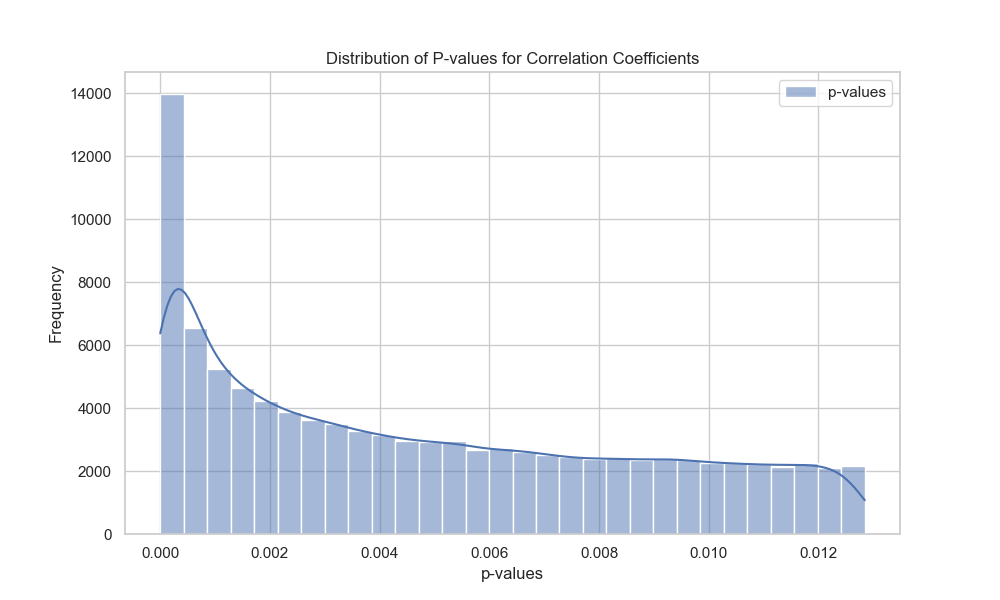

### Supplemental Figure 4

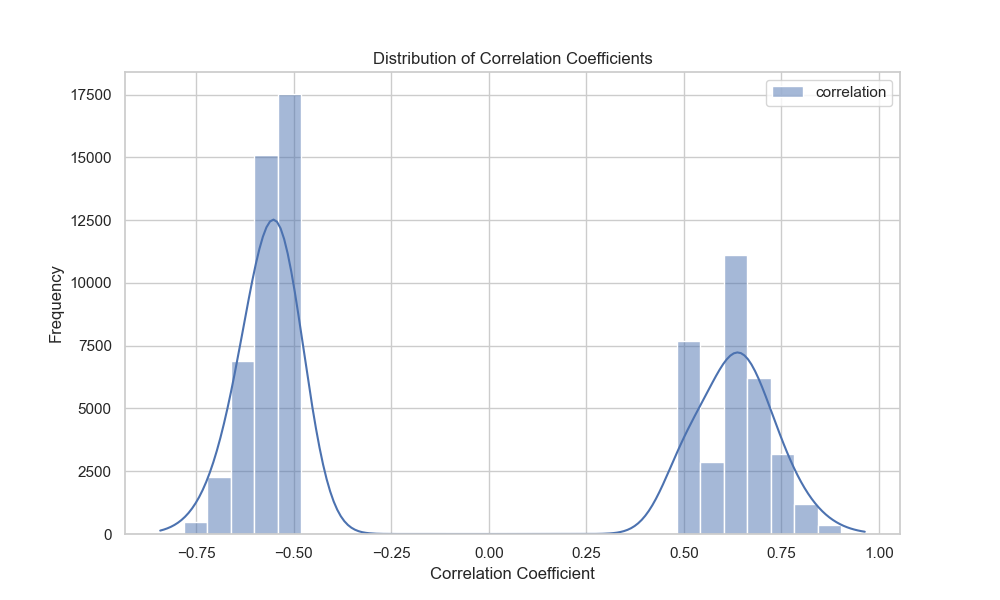

### Supplemental Figure 5

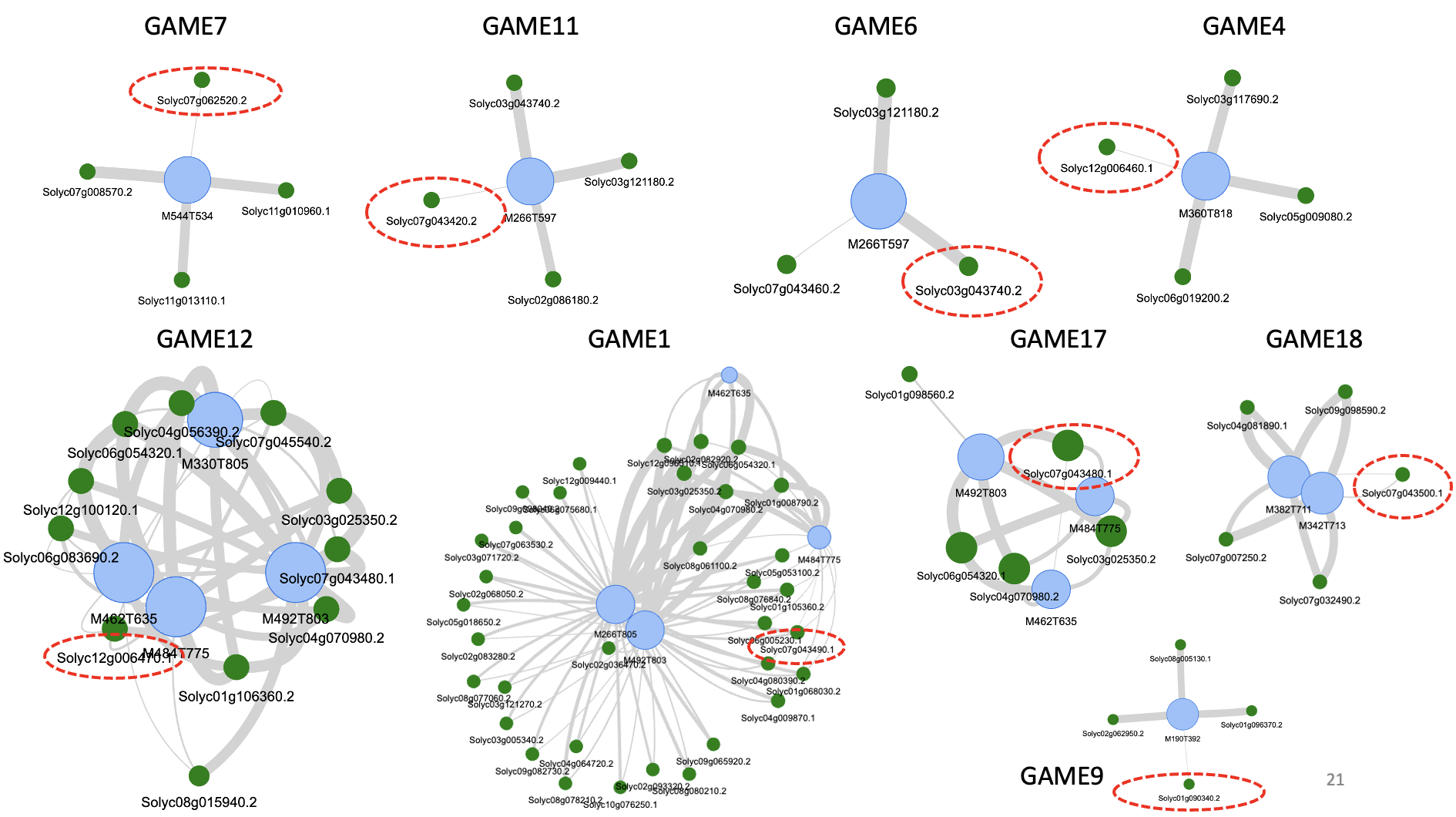

### Supplemental Figure 6

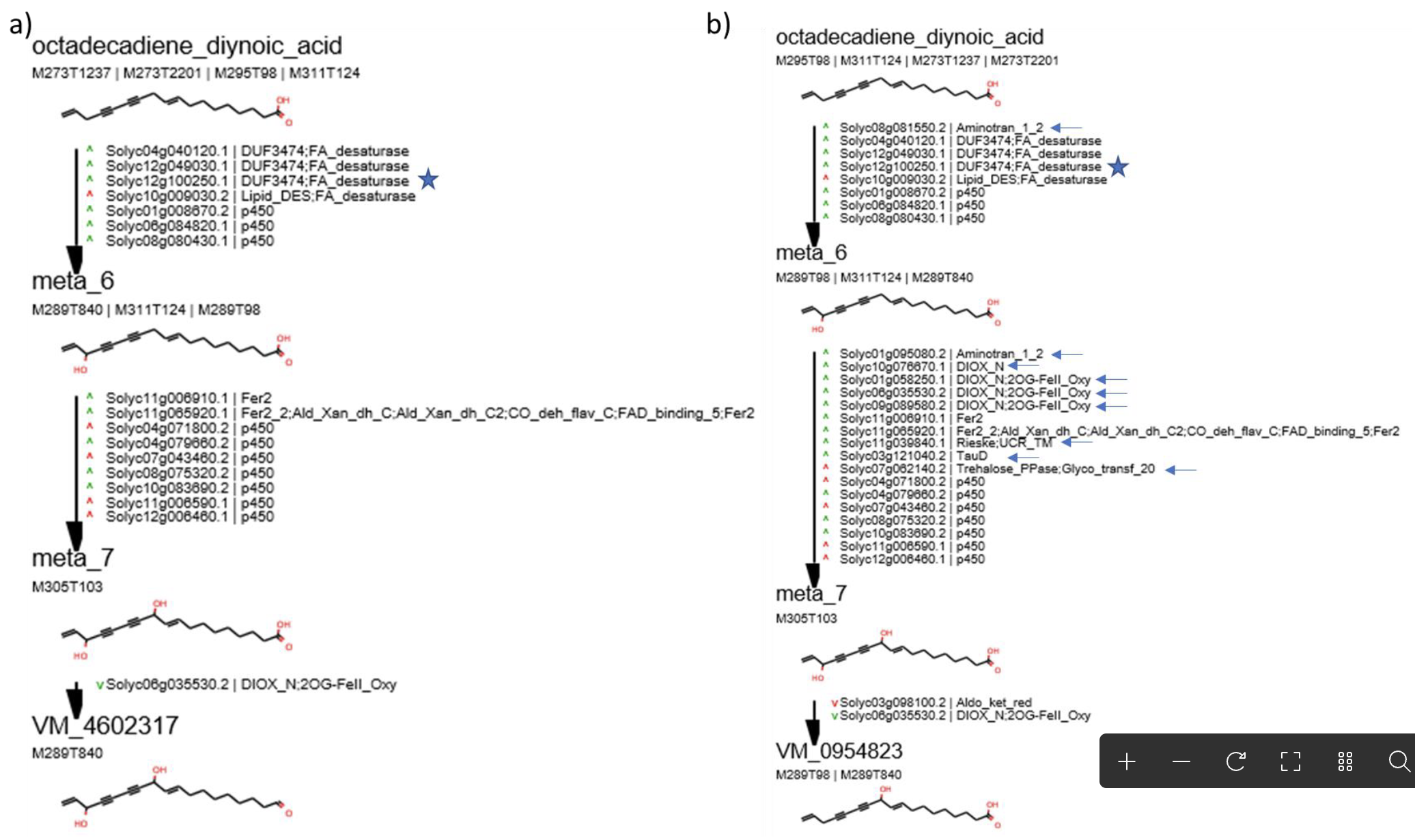

### Supplemental Figure 7

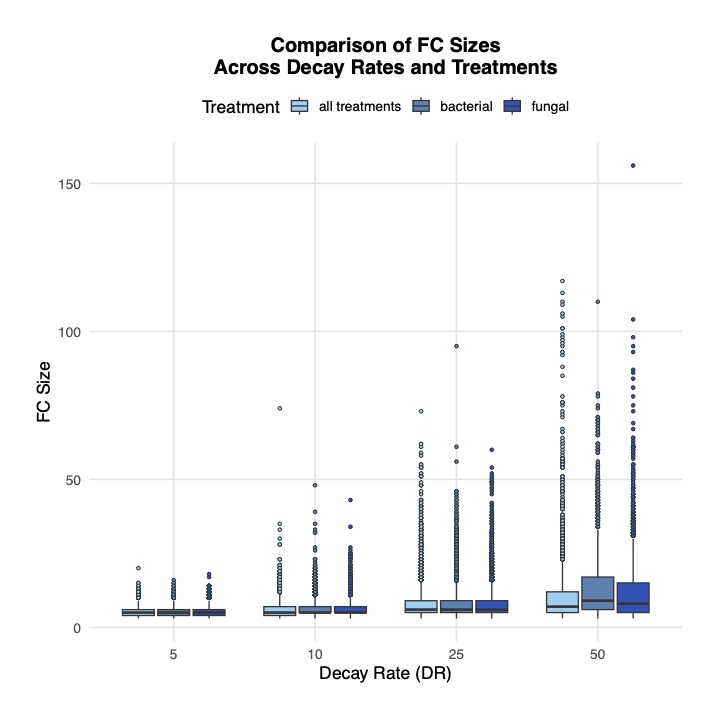

### Supplemental Figure 8

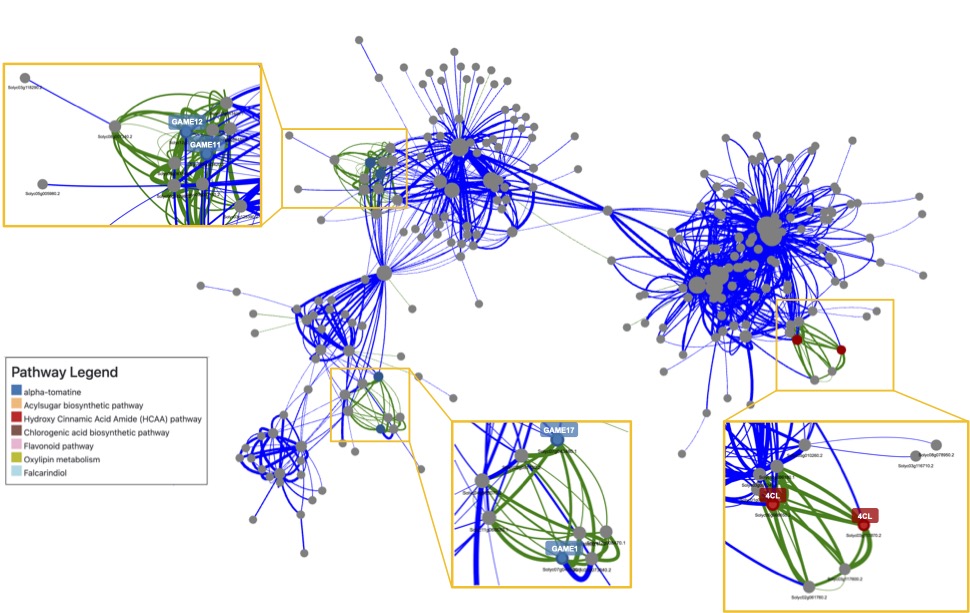
